# Myeloid Ecotropic Viral Integration Site 1 Promotes Vascular Smooth Muscle Differentiation through Activation of Myocardin

**DOI:** 10.64898/2026.09.26.751066

**Authors:** Alshimaa Wally, Sailesti Joshi, Jaser Doja, Ajay Kumar, Orazio J. Slivano, Susan H. Griffin, Kunzhe Dong, Ljubica Matic, Gabor Csanyi, Ulf Hedin, Xiaochun Long, Amr R. Salem

## Abstract

Myocardin (MYOCD) is the principal trigger-switch for vascular smooth muscle cell (VSMC) growth arrest and the cyto-contractile program of differentiation. While much is known about MYOCD-dependent VSMC differentiation, limited information exists as to its transcriptional regulation. Here, we report moderate mRNA expression of *Myeloid Ecotropic viral Integration Site 1* (*MEIS1*) in both human and mouse VSMCs. *MEIS1* mRNA is reduced in parallel with myocardin mRNA and protein under conditions of VSMC de-differentiation. Loss- and gain-of-*Meis1* expression studies in cultured human VSMCs and the mouse carotid artery support MEIS1 as an activator of myocardin and VSMC differentiation. CRISPR-mediated knockout of the endogenous *Meis1* gene *in vitro* and elicited lower levels of VSMC contractile markers and an increase in growth rate. Similarly, Femoral artery wire injury, in mice lacking *Meis1* showed and increased of neointima formation. Mechanistically, we identified a conserved MEIS1 response element in the proximal promoter of myocardin, that appears to contribute to the promoter activity, suggesting a potential regulatory interaction. These results suggest a previously unrecognized role for the homeodomain protein MEIS1 in the activation of myocardin and the quiescent, VSMC differentiated state.

## 1. Introduction

Under homeostatic conditions, vascular smooth muscle cells (VSMCs) exist in a condition of quiescence and contractile competency. Membrane-associated proteins and signaling pathways control this state of differentiation through the transcription of VSMC-restricted genes encoding cytoskeletal and contractile proteins ^1,2^. Several DNA-binding transcription factors carry out the task of activating this unique program of gene expression ^3^. One such factor is serum response factor (SRF) ^4^, which binds a 10-base pair *cis*-acting element known as a CArG box ^5^ located in the promoter or intronic region of a growing number of VSMC-restricted genes ^6,7^. As with most DNA-binding proteins, SRF is a weak activator of transcription. Consequently, SRF must interact with one of more than 70 cofactors to ensure proper levels of CArG-dependent gene expression necessary for the cell type or condition ^8^. One such SRF cofactor is Myocardin, first cloned in an *in silico* screen for cardiac-specific genes ^9^. Myocardin is a key component of a molecular switch for VSMC differentiation and quiescence ^10^, a finding confirmed in several subsequent reports ^11–13^.

Injury to the vessel wall provokes a limited number of resident VSMCs to de-differentiate, migrate out of the medial compartment towards the site of injury, and clonally expand ^14,15^. Combined lineage tracing and single cell transcriptomics clearly demonstrate phenotypic diversity of VSMCs in both human and experimental atherosclerosis ^16–18^. Attenuated expression of the myocardin cofactor ^19–22^ or loss of the SRF-Myocardin interaction over CArG boxes ^23–27^ facilitates the transition of VSMCs to a number of states. Such fate and state change likely depends on local environmental cues ^28^.

Studies of myocardin gene transcription have described functional *cis*-acting elements in the 5’ promoter region of myocardin that bind NKX2-5 ^29–33^, TP53 ^34^, FOXO3/4 ^35,36^, MYB ^37^, SRF ^38^, KLF4 ^39^, HNRNPA1 ^40^, and NFATC3 ^41^. These findings were limited to cell culture models and, with the exception of the NFATC3 binding site ^41^, the *cis*-acting elements were not conserved across multiple species suggesting species-specific promoter activation. The only in vivo interrogation of *myocardin* transcription to date used a lacZ reporter assay to define a functional enhancer located ∼30 kilobases upstream of the mouse *myocardin* transcription start site ^42^. Whether this enhancer is active in its resident genomic milieu has yet to be reported.

The CArGome comprises all hypothetical SRF-binding CArG boxes in the human and mouse genomes ^43^. In a bioinformatic screen for new CArG-dependent transcription factors, we discovered several CArG boxes around the *Myeloid Ecotropic viral Integration Site 1* (*MEIS1*) locus, including a conserved CArG-like box in the proximal promoter. Studies show that while MEIS1 is SRF and MYOCD-independent, VSMC de-differentiation results in its attenuated expression. Functional studies in cultured cells and the vessel wall show that MEIS1 induces Myocardin mRNA and protein expression and select contractile genes while suppressing growth. Importantly, a conserved MEIS1 response element in the proximal promoter of *myocardin* appears to bind MEIS1 and directly activate myocardin transcription.

## 2. Materials and methods

### 2.1 Cell culture transfection

The HCAMSC cells from Thermo Scientific (C0175C), Lonza (CC-2583), and Promo (C-12511) were cultured in Vasculife SMC Medium (LL-0014), and PAC1, A7r5 were cultured in DMEM combined with 10% Fetal Bovine Serum from Thermos Scientific (11965126, and10082147-respectively) at 37 °C with 5% CO2 incubator. Lipofectamine 3000 and Opti-MEM I from Thermo Scientific (L3000015 and 31985070-respectively) were used for transient transfection according to the manufacturer’s protocol.

### 2.2 Mouse aortic smooth muscle cell isolation

Primary mouse aortic smooth muscle cells (MASMCs) were isolated from pooled aortae (3-4 mice per preparation). Aortae were harvested into HBSS and digested for 20 min at 37°C in HBSS containing 1 mg/mL collagenase II (Worthington, LS004174) and 1 mg/mL soybean trypsin inhibitor (Worthington, LS003570). Following digestion, the adventitia was carefully removed, the aortae were opened longitudinally, and the endothelial layer was removed with a cotton swab. Aortae were minced and further digested for 40 min at 37°C in collagenase II/soybean trypsin inhibitor supplemented with elastase (Worthington, LS002279), with gentle trituration during digestion. Digestion was terminated with DMEM-based growth medium containing 20% FBS, 1% penicillin-streptomycin, and Smooth Muscle Growth Supplement (Thermo Fisher, S00725). Cells were centrifuged at 1,500 rpm for 5 min, resuspended in growth medium, and plated in 6-well plates. Primary MASMCs were expanded to confluence before subsequent experiments.

### 2.3 Generation of *Meis1*-knockout MOVAS cells

*Meis1*-null MOVAS cells were generated using CRISPR-Cas9 with two sgRNAs flanking the promoter and first coding exon of *Meis1*, designed to delete a 3.79-kb genomic region. sgRNA sequences and genotyping primers are listed in **Supplementary Table S5**. Following genome editing, single-cell clones were expanded by limiting dilution and screened by genotyping PCR. The expected deletion was confirmed by Sanger sequencing, and two independent *Meis1*-null clones were selected for subsequent experiments. The same sgRNA target sites were subsequently used for loxP knock-in during generation of the conditional *Meis1* mouse allele.

### 2.4 Western blotting

All steps were done according to the manufacturer’s protocol, using a Bio-Rad system (Bio-Rad, USA). Briefly, for mouse tissues were homogenized in RIPA lysis buffer (Sigma-Aldrich #R0278) using a Precellys CKMix tissue homogenizing kit, Bertin #P000918-LYSK0-A.0) with protease inhibitors cocktail (Sigma-Aldrich #04693159001), while cells were directly in protease inhibitor cocktail supplemented RIPA buffer, after that centrifugation at 12,000g at 4°C for 10 min. The supernatant was collected, and protein concentration was determined using BCA protein assay kits (Bio-Rad #500-0115, ##500-0114, and #500-0113), after that, protein samples were heated for 10 min at 95°C. Samples were run in a 4-20% Criterion TGX Stain-free SDS-PAGE (Bio-Rad #5678093). Samples were run in 4-20% SDSPAGE gel (Bio-Rad, 4561094) in 1x Tris-Glycine running buffer (Bio-Rad, A0028), after that, proteins were transferred to PVDF membrane (Bio-Rad #1620175) using Trans-blot Turbo transfer. Membranes were blocked in a blocking buffer (Bio-Rad, 12010020) and incubated with appropriate antibodies (Supplementary Table 1). Bands were visualized by enhanced chemiluminescence method (ECL) Advasta (#K-12043-D20) and ChemiDoc imaging system (Bio-Rad, 12003154). The protein bands intensity was quantified using Image J program (National Institutes of Health, Bethesda, MD, USA).

### 2.4 Viral vectors

The lenti-viral vectors both scrambled and Meis1were customized vectorBuilder (Meis1 shRNA1 # LVM(VB201217-1137wmv), Meis1 shRNA2 # LVM(VB201217-1139vhs), and shRNA scrambled# LVM(VB010000-0001mty)-C) and adenoviral vectors for in vivo transduction was customized from Vector Biolabs (shAAV-264485 for knockdown and AAV26445 for overexpression studies), adeno-GFP (#1060) was used as a control. For Cre-mediated recombination of the floxed *Meis1* allele, Cre-EGFP adenovirus (VectorBuilder; VB260327-1359uxf) was used. In vitro viral transduction was done using polybrene EMD Millipore (#TR-1003-G) and 3-5 viral transduction unit/cell. GFP was used as a transduction control.

### 2.5 Real Time PCR

Total RNA was extracted from cells and tissues using RNeasy Plus Mini Kit (Qiagen #74134) or TRIzol (Thermo Fisher #15-596-018) following the manufacturers’ instructions. TRIol extracted RNA, was treated with RQ1 RNase-Free DNase (Promega #M6101) to eliminate genomic DNA contamination. Reverse transcription was performed using Iscript Reverse Tranascription Supermix, (Bio-Ra#1708841), and RT-qPCR was performed in a 10µl and SYBR Green Master Mix (Bio-Rad #1708882) on a Bio-Rad detection instrument (CFX Connect). List of RT-qPCR primers were tabulated in **Supplementary table 1.**

### 2.5 Immuno- RNA *in situ* hybridization

Human coronary atherosclerosis samples were obtained from Dr, Csanyi, and for mice coronary artery were fixed in 10% neutral buffered formaldehyde, paraffin embedded, and sectioned at 5μm. Sectioned were processed for a combined immunofluorescence of TAGLN (abcam# ab14106), a marker for smooth muscle, and RNA in situ hybridization of Meis1 mRNA (#436361, ACD) according to the manufactures instructions. Alexa fluor 488 secondary antibody was used to detect TAGLN protein (Thermo Fisher). Signals were obtained with a LSM 900 confocal laser scanning microscope (Zeiss) using the Zeiss Blue software system for image acquisition and processing.

### 2.6 Pluronic Gel

Pluronic gel was prepared as described previously ^44^. Briefly, 25% Pluronic F127 (Sigma-Aldrich #P2443), 2% Polycarbophil (PCB) (Lubrizol), and 1% Trypsin (Sigma-Aldrich #T4799) were dissolved in 1xPBS, kept on rotation overnight at 4°C. 4 x10^8^ of viral particles were dissolved in 50ul of Pluronic gel, and wrapped around the left carotid artery of 3xHA-*Myocd* mouse model ^45^ for two weeks after that, LCA was isolated for RNA, Western blotting and Immunostaining.

### 2.7 arterial denudation and injury models

To evaluate the effects of meis1 deletion in vascular remodeling, *Meis1*^fl/fl^-*Itga8Cre*ER^T2^ mice (KO) and *Meis1*^wt^-*Itga8Cre*ER^T2^ (Ctrl) (10 weeks old) were administered tamoxifen (50 mg/kg, ip) once daily for 5 consecutive days. After a two-week washout period, mice underwent wire injury of the left femoral artery and complete ligation of the left carotid artery, as previously described^46,47^. Following injury, mice were maintained for 4 weeks before euthanasia. The injured femoral and carotid arteries and their contralateral uninjured counterparts, used as internal controls, were then harvested, fixed in 4% paraformaldehyde and subjected to histological analysis.

### 2.8 BiKE and BiRCA transcriptomic analyses

Human carotid transcriptomic data were obtained from the Biobank of Karolinska Endarterectomy (BiKE)^48^, which includes carotid endarterectomy specimens and associated clinical data from patients with carotid atherosclerosis^49^. MEIS1 expression was compared between control arteries and carotid plaques and between asymptomatic and symptomatic plaques. The Biobank of Rat Carotid Artery injury (BiRCA) ^50^dataset was used to assess temporal changes in Meis1 expression following carotid balloon injury, with injured and contralateral arteries analyzed across multiple post-injury time points.

### 2.9 Statistical analysis

Each in vitro experiment was repeated at least two times and at least in two different smooth muscle cells. Statistical significance among the groups were analyzed with one-way ANOVA followed by a post-hoc Tukey’s test. For two groups analysis, an independent t-test was used for statistical analysis. The statistical significance was established at 0.05 (*, P≤0.05, ** P≤0.01, ***, P≤0.001) using SPSS® software V.21 (Chicago, IL), and GraphPad Prism 9.0 used for illustrations (GraphPad Software, La Jolla, CA, USA).

## 3. Results

### 3.1 MEIS1 is a CArG-MYOCD-SRF independent transcription factor in VSMCs

The 5’ *MEIS1* locus harbors several conserved CArG boxes, including a non-consensus CArG box (CCTGTTATGG) in the promoter region. This proximal CArG box exhibits SRF-binding as shown by ChIP-seq of human coronary artery SMCs (HCASMC) and human umbilical vein endothelial cells as well as several cell types from ENCODE (**Supplementary Fig. 1A**). Luciferase assays revealed mild SRF-dependent activation of this proximal CArG box (**Supplementary Fig. 1A**). However, lentiviral-mediated MYOCD loss- and gain-of-function studies failed to show any change in mRNA expression of *MEIS1* or that of its two paralogs, *MEIS2* and *MEIS3* (**Supplementary Fig. 1B, 1C**). Knockdown of SRF in three different lots of HCASMC (Lonza, Promo, and Thermo) demonstrated no change in steady-state mRNA expression of *MEIS1*, *MEIS2*, and *MEIS3* (**Supplementary Fig. 1D, 1E** and **Supplementary Fig. 2A, 2B**). We next performed bulk RNA-seq following lentiviral-mediated knockdown of SRF, which resulted in a more than 90% reduction of *SRF* mRNA in each lot of HCASMC (**Supplementary Fig. 2C**). Consistent with the above findings, there was no change in *MEIS1* mRNA across each HCASMC lot (**Supplementary Fig. 2D**), although an expected reduction was seen in known SRF targets such as *TAGLN* and *CNN1* (**Supplementary Fig. 2E, 2F**). Interestingly, we also noted no change in *MYOCD* mRNA with SRF knockdown (**Supplementary Fig. 2F**). These results indicate that despite the presence of conserved CArG boxes, *MEIS1* mRNA expression is independent of the SRF-MYOCD transcriptional switch.

### 3.2 *MEIS1* expression is reduced under conditions of VSMC de-differentiation

Encouraged by the moderate expression levels of *MEIS1* mRNA in HCASMC, as well as a previous report showing altered MEIS1 in pulmonary artery hypertension ^51^, we carried out additional expression studies in VSMC undergoing de-differentiation. Because no reliable antibody exists to MEIS1 (**Supplementary Fig. 5**), we performed RNA-scope on sections of human atherosclerotic coronary artery (**Fig. 1A**). These results demonstrated easily detectable *MEIS1* mRNA in the medial layer, but much lower levels in the atheroma (**Fig. 1B**) where de-differentiated VSMCs undergo cell state changes ^16–18^. The expression changes seen with *MEIS1* paralleled those of MYH11 protein (**Fig. 1B**). In line with our findings, transcriptomic analysis from <u>Bi</u>obank of <u>K</u>arolinska <u>E</u>ndarterectomy (BiKE)^48^ showed a significant down-regulation of *MEIS1* mRNA expression in both plaques compared to control arteries (**Fig 2A**), and in plaques from symptomatic versus asymptomatic patients (**Fig 2A**). Additionally, snATAC-Seq analysis of human carotid atherosclerosis^52^ showed *MEIS1* is mainly accessible in VSMC cell cluster (**Fig 2C, 2D**), and it’s accessibly decreased in modulated VSMC (**Fig 2D**). A spatiotemporal transcriptomic profiling of the rat left carotid balloon injury, a rodent model of intimal hyperplasia, showed a significant reduction of *Meis1* as early as 2hrs post injury (**Fig 2E**), a time point which is earlier than the classical contractile markers shown from 20hrs to 5days (middle phase)^50^(**Fig 2E**). Of note, the reacquisition of *Meis1* expression in the later stages was associated with restoration of the contractile features, suggesting *Meis1* is an upstream activator of classic differentiation markers. Additionally, the middle phase was associated with tissue remodeling, SMC proliferation, and migration strongly suggesting that reduced MEIS1 is a potent driver of SMC phenotypic modulation

**Figure 1.**
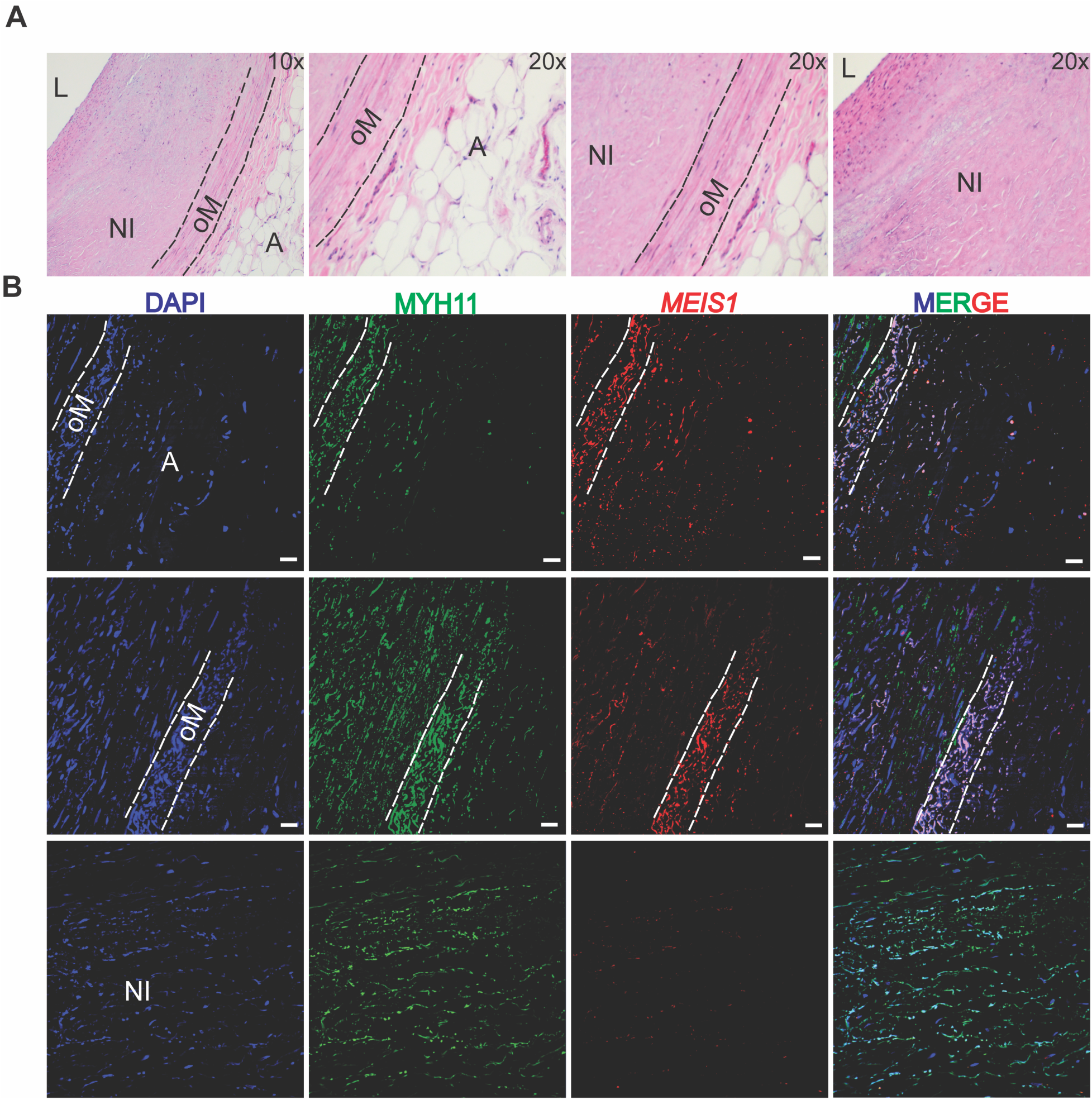
*MEIS1* is down regulated in human coronary atherosclerosis. **A)** Hematoxylin and Eosin staining of atherosclerotic human coronary artery (A= adventitia, oM= Outer VSMC layer, NI= neointimal, and L= Lumen). B) *MEIS1* RNA-scope combined with immunostaining of MYH11. Scale bar is 40x.

**Figure 2.**
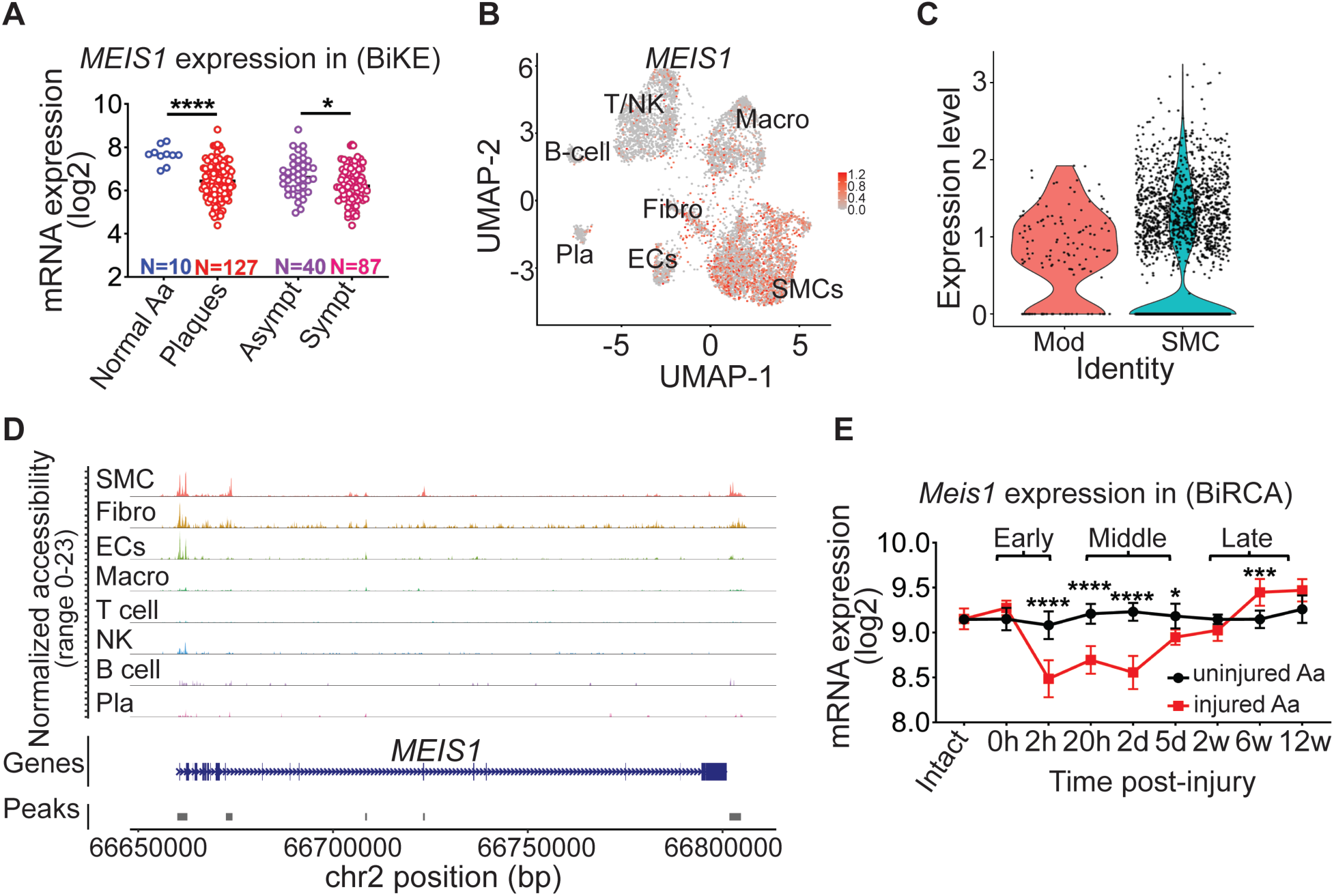
*MEIS1* expression, accessibility in human atherosclerosis, and rat carotid artery injury. **A)** *MEIS1* expression in normal arteries, carotid plaques from symptomatic and asymptomatic patients. **B**) UMAP plot of *MEIS1* in different atherosclerotic cell sub-clusters. **C**) Violin plot of MEIS1 distribution in contractile (SMC) and modulated (mod) SMCs clusters data from (**B**). **D**) MEIS1 cell type specific chromatin landscape. **E**) Spatiotemporal expression of *Meis1* after carotid balloon-injury in rats (BiRCA)

GTEx portal expression data in human tissues show highest expression of *MEIS1* mRNA in SMC-rich tissues such as uterus and gastrointestinal organs with lower levels in aorta and coronary artery. A similar profile is seen with *MEIS2* and MEIS3 (**Supplementary Fig. 3A-4C**). We next investigated *Meis1* mRNA expression in the mouse. Both male and female mice exhibited moderate levels of *Meis1* mRNA in aorta with vanishingly low levels of *Meis2* and *Meis3* (**Supplementary Fig. 3D-3F**). Among the three *Meis* paralogs, *Meis1* showed the highest expression levels in normal mouse aorta (**Fig. 3A**). As expected, passage of mouse aortic SMCs (MASMCs) resulted in decreased expression of *Myocd*, *Myh11*, and *Cnn1* mRNA whereas the same cultures showed a stepwise increase in *Klf4* mRNA (**Fig. 3B**). MASMCs derived from an HA-tagged *Myocd* mouse ^45^ showed parallel reductions in MYOCD and MYH11 protein (**Fig. 3C**). Of note, *Meis1*, but not *Meis2* or *Meis3*, displayed attenuated mRNA expression in passaged MASMCs (**Fig. 3D**). Next, we interrogated mRNA expression of the same gene set in the ligated mouse carotid artery, which is a model of VSMC de-differentiation. Levels of *Myocd*, *Myh11*, and *Lmod1* mRNA were diminished in the injured carotid artery, with expected increases in *Ki67* mRNA (**Fig. 3E**). Interestingly, expression of both *Meis1* and *Meis2* mRNA was reduced in the injured carotid artery, and *Meis3* showed a downward trend (**Fig. 3F**). To probe the expression of *Meis1* mRNA further, we used RNA-scope and observed expression in the medial layer, but no detectable signal in the neointima (**Figure 3G**). Taken together, these findings demonstrate attenuated *Meis1* mRNA in several conditions VSMC de-differentiation.

**Figure 3.**
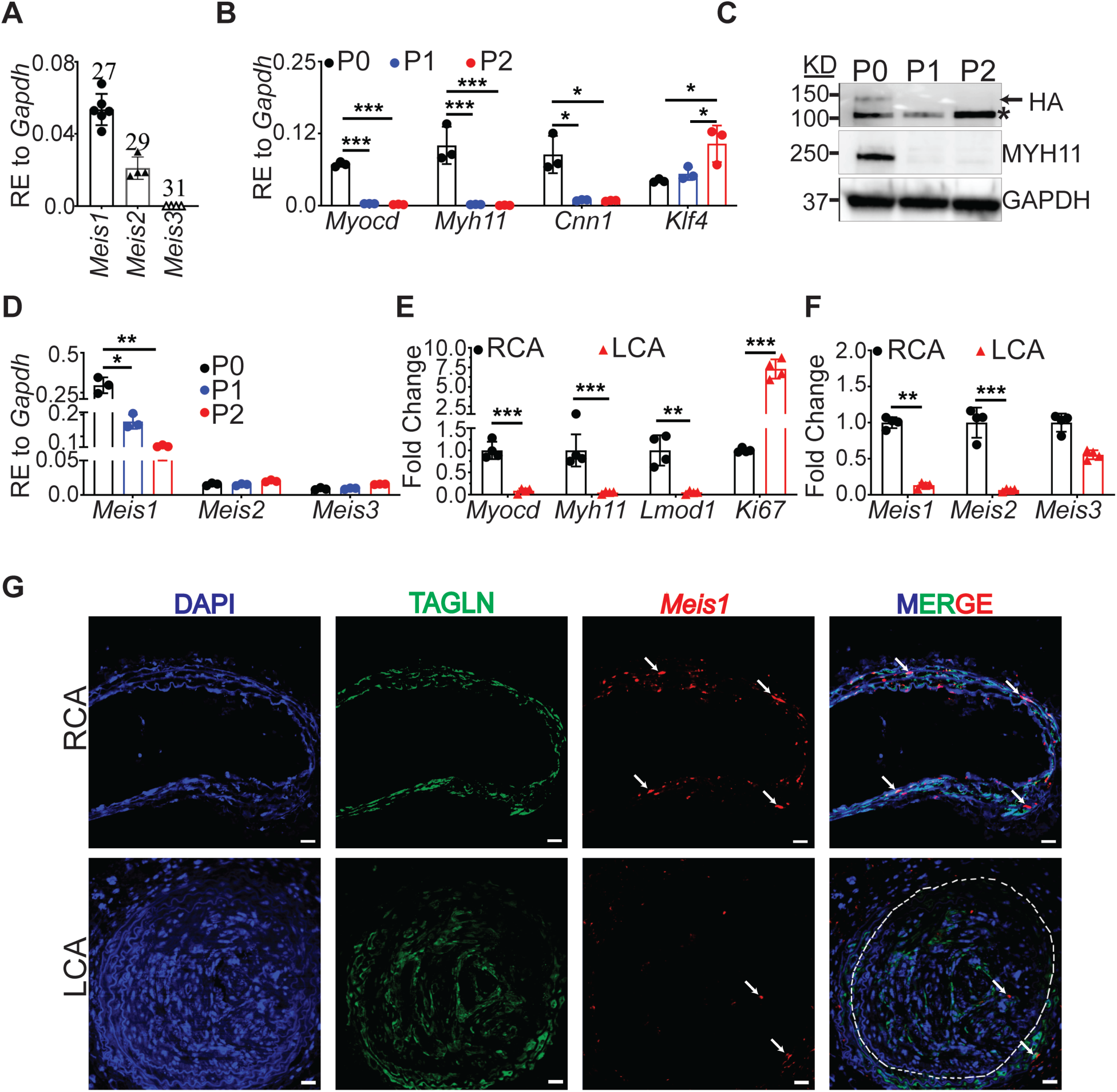
*Meis1* is down regulated in vitro and in arterial injury models. **A)** qRT-PCR of *Meis1*, *Meis2*, and *Meis3* in mouse aortae. **B**) qRT-PCR of *Myocd*, *Myh11*, *Cnn1*, and *Klf4* in <u>M</u>ouse <u>A</u>ortic <u>S</u>mooth <u>M</u>uscle <u>C</u>ell (MASMC) serial passages (P0, P1, P2). **C**) qRT-PCR of *Meis1*, *Meis2*, and *Meis3* in MASMC serial passages. **C**) Western-blotting of HA-MYOCD, and MYH11 in MASMC serial passages (Arrow= HA, * non-specific). **D**) qRT-PCR of *Myocd*, *Myh11*, *Lmod1*, and *Ki67* in mouse carotid arteries (RCA= un-ligated <u>R</u>ight <u>C</u>arotid <u>A</u>rtery, LCA= ligated <u>L</u>eft <u>C</u>arotid <u>A</u>rtery). **E)** qRT-PCR of *Meis1*, *Mesi2*, and *Mesi3* in RCA and LCA. **F**) *Meis1* RNA-Scope combined with TAGLN immunofluorescence in un-ligated RCA, and Ligated LCA (White arrows), a representative image of three biological repeats. Each dot represent the mean of three technical replicates.

**Figure 4.**
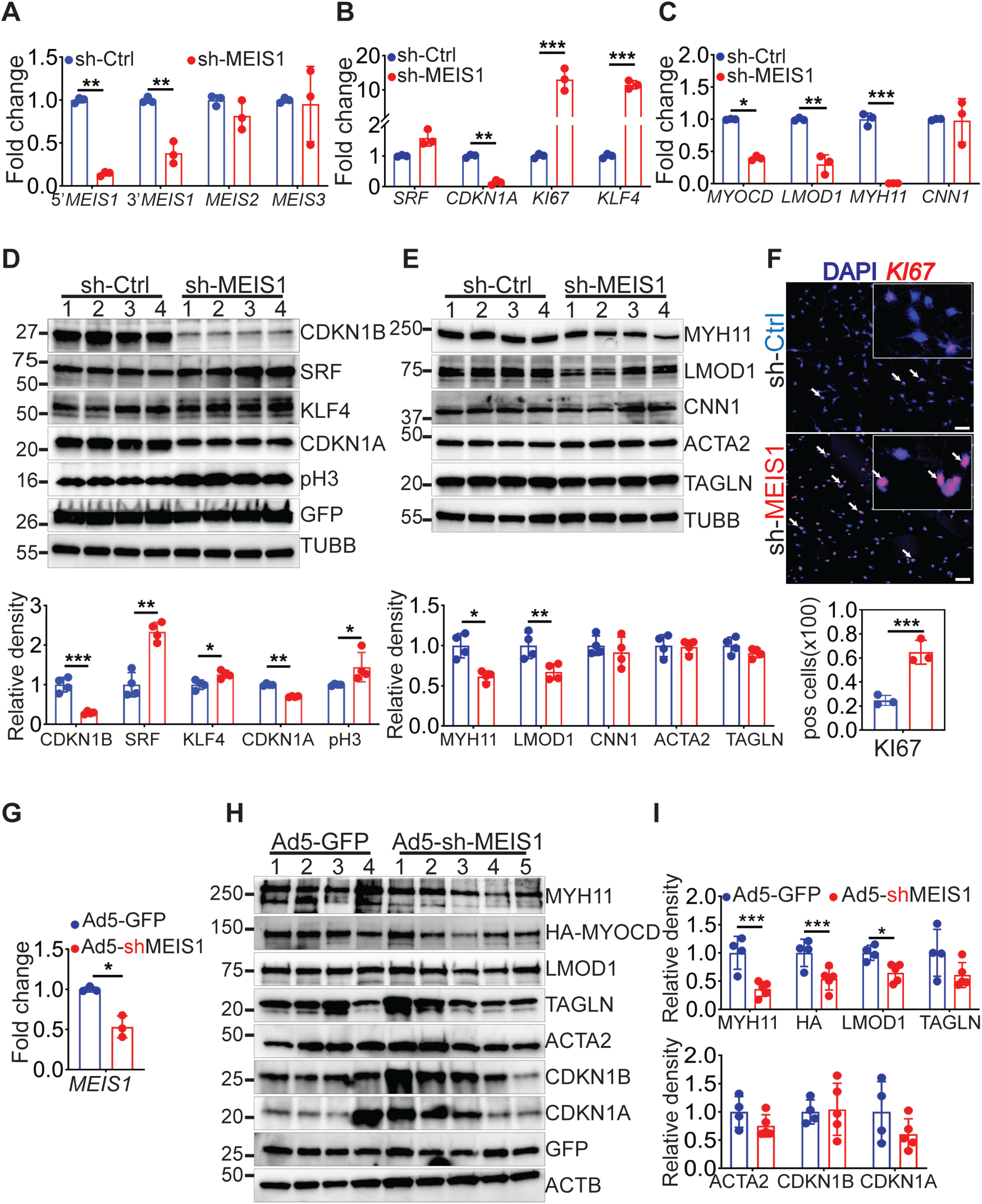
Loss of *Meis1 in vitro* and *in vivo* increases VSMC de-differentiation. **A)** qRT-PCR of *MEIS1*, *MEIS2*, and *MEIS3* in HCASMC (sh-ctrl= scrambled shRNA, and sh-MEIS1= shRNA specific to *MEIS1*). **B**) qRT-PCR of *MYOCD*, *LMOD1*, *MYH11*, *CNN1* in *MEIS1* deficient HCASMC. **C**) qRT-PCR of *SRF*, *CDKN1A*, *KI67*, *KLF4* in *MEIS1* deficient HCASMC. **C-D**) Western blotting of Proliferation markers (CDKN1B, KLF4, CDKN1A, pH3), and contractile markers (MYH11, LMOD1, CNN1, ACTA2) respectively below is the quantification (GFP is a transduction control and TUBB is a loading control). **F**) Immunostaining of KI67 in MESI1 knockdown HCASMC and below is the quantification. **G-I**) **Ectopic in vivo loss of *Meis1***, G) qRT-PCR of *Meis1*. **H-I**) Western blotting of contractile markers (MYH11, HA-MYOCD, LMOD1, TAGLN, ACTA2), and proliferation markers (CDKN1A, CDKN1B) (GFP is adenovirus transduction control, and TUBB is and endogenous loading control), quantification in (**I**).

### 3.3 *MEIS1* knockdown elicits VSMC de-differentiation in vitro and in vivo

A lentivirus engineered to carry a short-hairpin RNA to *Meis1* was used to interrogate VSMC phenotype in HCASMCs. Effective and specific knockdown of *MEIS1* mRNA was observed in HCASMCs transduced with the shRNA to *MEIS1* but not a control shRNA with scrambled sequence (**Fig. 4A**). *MEIS1* knockdown resulted in attenuated mRNA expression of the cell cycle inhibitor *CDKN1A* and an increased level of *KI67* and *KLF4* mRNA (**Fig. 4B**). HCASMC transduced with the shRNA to *MEIS1* showed attenuated expression of *MYOCD*, *LMOD1*, and *MYH11* mRNA (**Fig. 4C**). Comparable changes in protein expression were also observed with MEIS1 knockdown (**Fig. 4D, 4E**). Of note, and consistent with increased phospho-histone 3 (pH3) protein expression (**Fig. 4D**), knockdown of *MEIS1* stimulated HCASMC DNA synthesis (**Fig. 4F**). We next investigated the effects of loss of *Meis1* expression in the intact carotid artery via pluronic gel-mediated release of Adenoviral-shuttled shRNA to *Meis1* (**Supplementary Fig. 3A**). A GFP tag confirmed effective targeting of adventitial and medial cells of the carotid artery 14 days post-transduction (**Supplementary Fig. 3B**). qRT-PCR of the carotid artery revealed a ∼50% decrease in *Meis1* mRNA with viral-mediated knockdown of Meis1 (Fig. 3G). The decrease in Meis1 mRNA was associated with a diminution in VSMC contractile protein expression (MYOCD, MY11, LMOD1) and the cell cycle inhibitor CDKN1A (**Figure 4H-4I**). We also observed a qualitative decrease in ACTA2 staining of the carotid artery (**Supplementary Fig. 3B**).

### 3.4 *MEIS1* overexpression promotes a VSMC contractile state

We next utilized an adenovirus containing a GFP reporter and an expression sequence for *MEIS1* to transduce HCASMCs. Results showed an expected 200-fold increase in *MEIS1* mRNA expression with adenoviral delivery of *MEIS1* (**Fig. 5A**). A concomitant elevation in cell cycle inhibitors, CDKN1A and CDKN1B and contractile marker proteins (MYH11, LMOD1, CNN1, and ACTA2) was also observed indicating an advanced VSMC differentiated state (**Fig. 5B, 5C**). Next, pluronic gel-encapsulated adenovirus containing a *Meis1* expression cassette was applied to the left carotid artery. Here, a nearly 3-fold increase in *Meis1* mRNA was seen in experimental mouse carotid artery (**Fig. 5D**). This elevation in *Meis1* mRNA was accompanied by a modest increase in cell cycle inhibitor proteins and the cytoskeletal LMOD1 (**Fig. 5E, 5F**). A notable finding was an increase in the MYOCD cofactor, detected in a CRISPR-tagged mouse ^45^ using an HA antibody (**Fig. 5E, 5F**). These findings suggest that gain of Meis1 expression promotes, at least partially, a VSMC differentiated state.

**Figure 5.**
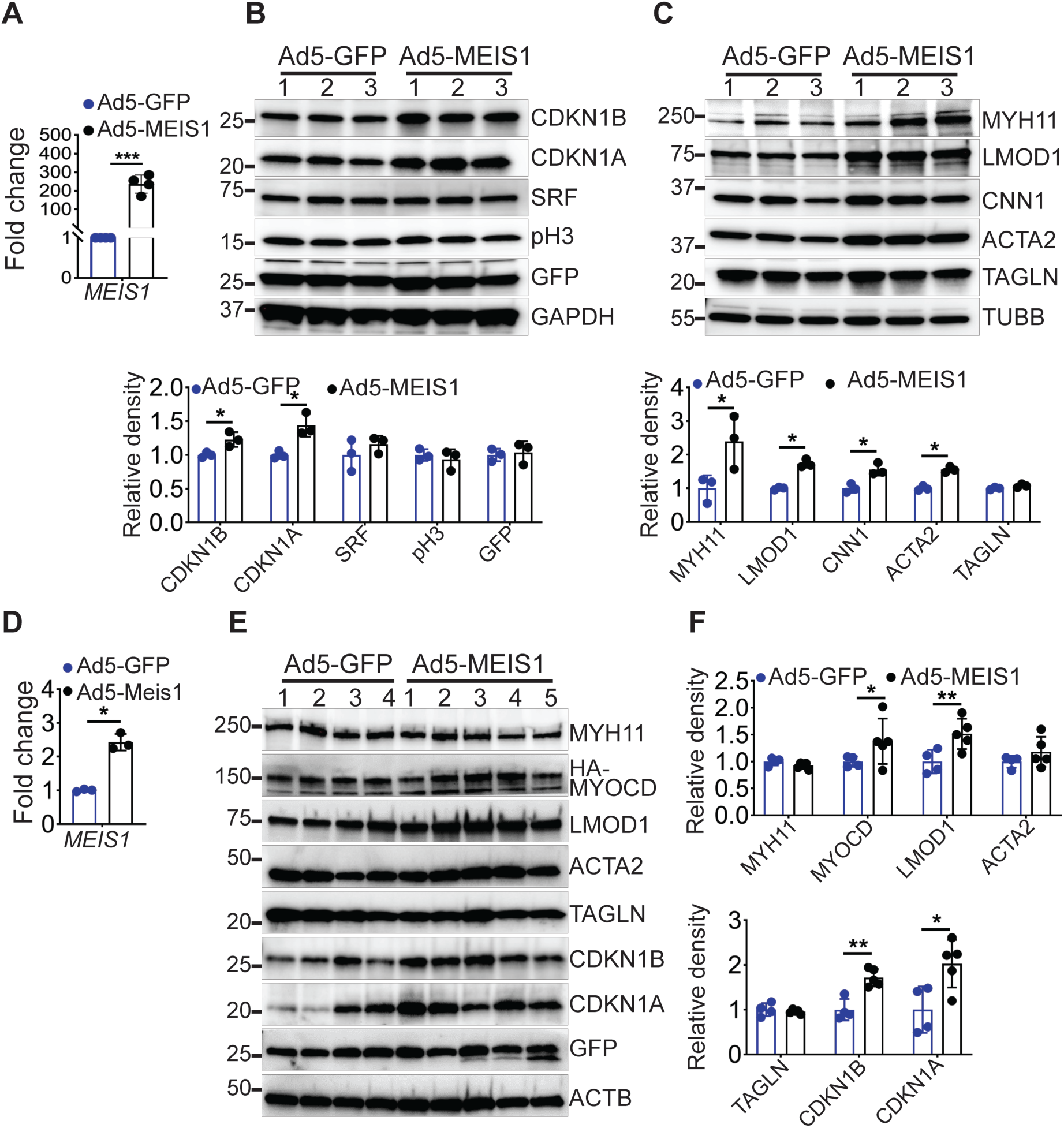
Gain of MEIS1 *in vitro* and *in vivo* maintains VSMC contractile state. **A)** qRT-PCR of *MEIS1* in HCASMC. **B**-**C**) western blot analysis of proliferation markers (CDKN1A, CDKN1B) and quantification below in (**B**), and contractile markers (MYOCD, MYH11, LMOD1, CNN1, ACTA2, TAGLN) and quantification below in (**C**). **D-F**) **ectopic delivery of MEIS1 through pluronic gel to the left carotid artery. D**) qRT-PCR of *Meis1*. **D-F**) western blotting of contractile (MYH11, MYOCD, LMOD1, ACTA2, TAGLAN), and proliferation markers (CDKN1A, CDKN1B). Western blot quantification in (**F**). 1,2,3,4, 5 are independent mouse, GFP used as a transduction control, and ACTB is a loading control.

### 3.5 CRISPR deletion of *Meis1* in an immortal VSMC line

The results thus far used ectopic loss- and gain-of-*Meis1* expression studies, which are limited by potential off-targeting events (with shRNA) and supra-physiological levels of *Meis1* (with adenoviral overexpression). We therefore used CRISPR-Cas9 gene editing ^53,54^, to inactivate the endogenous *Meis1* gene in an immortalized mouse aortic smooth muscle cell line (MOVAS), ^55^). As a previous knockout strategy resulted in an incomplete knockout (see Discussion), we developed a strategy that should result in a true null allele by targeting the promoter and first coding exon of the *Meis1* locus (**Fig. 6A**). PCR and Sanger sequencing of genomic DNA from two independent clones confirmed the 3.79 kilobase deletion (**Fig. 6B**) and qRT-PCR revealed barely detectable levels of *Meis1* mRNA (**Fig. 6C**). Western blot analysis showed a significant decrease of contractile markers (LMOD1, CNN1) and CDKN1A (**Fig. 6D**). Although the decrease in CDKN1A protein mirrored a similar decrease in *Cdkn1a* mRNA, *Lmod1* and *Cnn1* transcripts were, for unclear reasons, increased in the deletion clone and levels of *Myocd* were undetectable (not shown). On the other hand, cell proliferation was elevated in two independent *Meis1* null cell lines (**Fig. 6E**). Since, a number of commercial antibodies failed to detect the endogenous MEIS1 protein (**Supplementary Fig. 5A, 5B**), a 3x-HA N-terminal tag was tested in ectopic expression studies. Western blotting of cells co-transfected with an siRNA to *Meis1* and either of two HA-tagged *Meis1* isoforms confirmed detection of both isoforms at the correct molecular weights (**Fig. 5F**). Further, immunostaining of cells revealed nuclear localization of the HA-tagged MEIS1 protein suggesting the presence of the tag does not alter localization (**Figure 6G**). Building on these findings, we used the GFP-HA–tagged Meis1 mouse line from The Jackson Laboratory and introduced loxP sites flanking the same region targeted in our CRISPR deletion strategy, generating a conditional *Meis1* allele. Genotyping and recombination PCR confirmed the engineered allele, while HA immunostaining demonstrated nuclear localization of HAMEIS1 in medial VSMCs (**Fig. 6I, left**), which was markedly reduced following carotid artery ligation (**Fig. 6I, right**). Moreover, primary mouse aortic SMCs isolated from these mice and transduced with Cre virus showed complete loss of MEIS1 protein, validating efficient conditional deletion of the floxed allele (**Fig. 6J-K**). Together, these findings validate a true conditional null *Meis1* allele that enables reliable detection of endogenous MEIS1 and overcomes the incomplete knockout associated with the previously described exon 8-targeted floxed model (data not shown).

**Figure 6.**
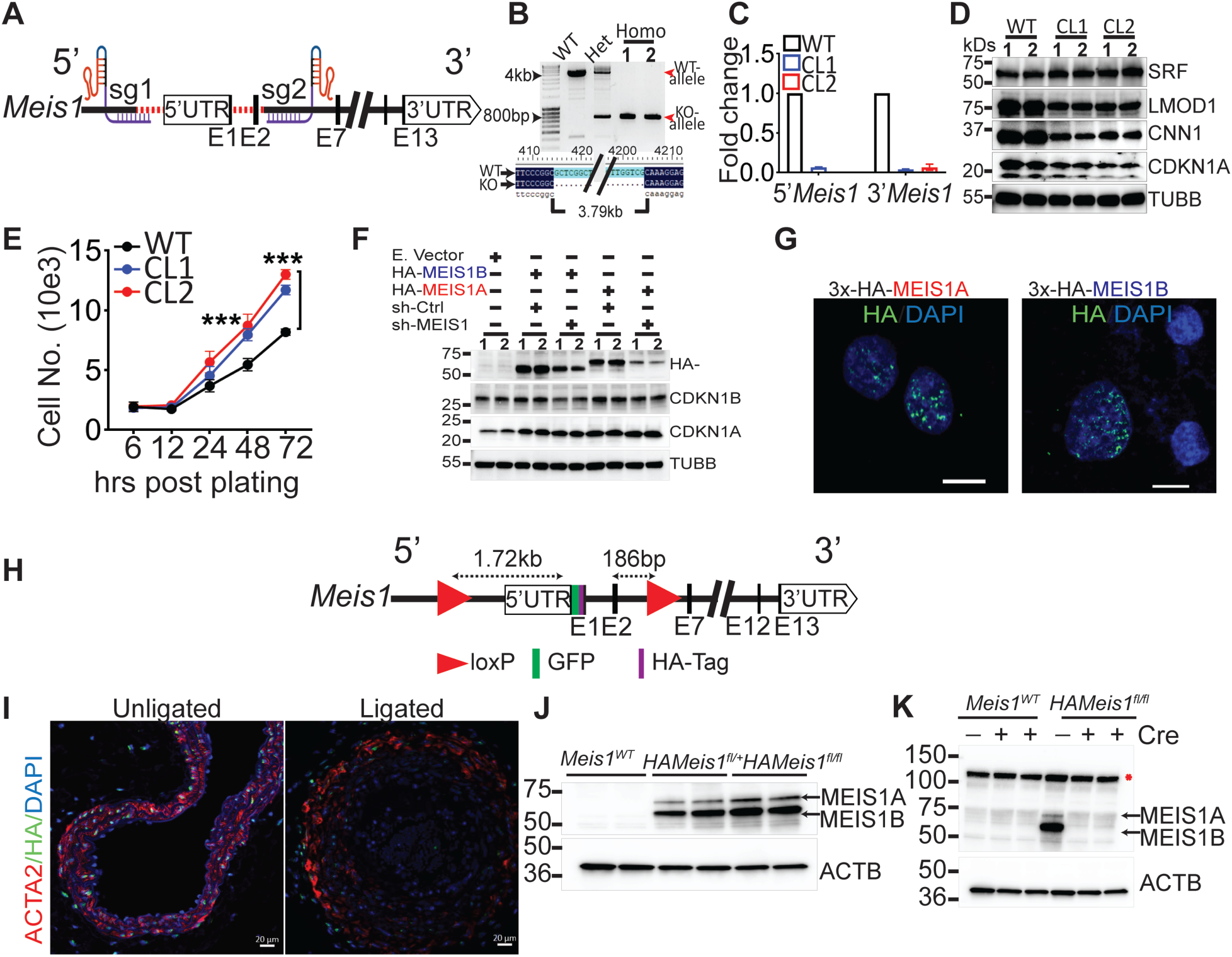
Strategic generation of *Meis1*Δ Cell line and floxed-tagged *Meis1* mice. **A)** Schematic representation of Meis1 locus indicating the positions of sgRNAs (sg1, sg2) used to generate a null *Meis1* mouse smooth muscle cells (MOVAS), and Exons (E1, E2,E3, E11,E12). **B**) Genotyping PCR showing WT-allele and KO-allele and Sanger sequencing confirmed a 3.79 kb deletion (below). **C**) qRT-PCR showed complete absence of *Meis1* 3’- and 5’-terminal (CL1, CL2= homozygous clones from (**B**). **D**) Western blot analysis of LMOD1, CNN1, CDKN1A. 1, 2 are duplicate loading. **E**) Cell proliferation assay in Meis1 null clones CL1, CL2. **F**) 3x-HA-tagge MEIS1A/MEIS1B in PAC1 SMC± sh-Meis1 or sh-random with decreases in HA, CDKN1B, and CDKN1A. **G**) Immunofluorescence of 3x-HA-N terminal tagged Meis1A, and Meis1B in rat aortic smooth muscle cells (PAC1). **H**) **Schematic representation of the floxed *HAMeis*1 allele**, with loxP sites flanking the targeted region. **I**) Immunofluorescence of ligated and unligated carotid arteries showing HAMEIS1 in ACTA2^+^ VSMCs. **J**) Western blot showing endogenous MEIS1A and MEIS1B protein in WT, heterozygous, and homozygous *HAMeis1* mice. **K**) Western blot of primary MASMCs isolated from WT and homozygous floxed HAMeis1 mice ± Cre, showing complete loss of MEIS1 protein following Cre-mediated excision. Red asterisk, nonspecific band

### 3.6 MEIS1 activates the *Myocd* promoter in vitro

The altered expression of *Myocd* mRNA and MYOCD protein with loss- or gain-of-Meis1 expression, respectively, suggested to us that MEIS1 could directly activate the *Myocd* promoter. To address this idea, we first interrogated the *Myocd* promoter region using the JASPAR database ^56^. This analysis revealed a consensus-binding site for the MEIS1 transcription factor located on the negative strand 75 base pairs upstream of the mouse *Myocd* transcription start site (**Fig. 7A**). The MEIS1 response element (MRE) is conserved across multiple mammalian species and is identical to an MRE located in the 5’ regulatory region of the known MEIS1 response gene, Cdkn1a (**Fig. 7B**). Luciferase assays showed an increase in *Myocd* and *Cdkn1a* promoter activity following co-transfection with either MEIS1A or MEIS1B isoforms (**Fig. 7C, 7D**). Such activation was abolished upon deletion of each MRE (**Fig. 7E, 7F**). ChIP-PCR of VSMC transfected with a FLAG-tagged *Meis1* expression cassette showed enriched DNA encompassing the MRE (Fig. 7G). Together, these findings suggest MEIS1 directly activates the transcription of *Myocd* through binding of an MRE.

**Figure 7.**
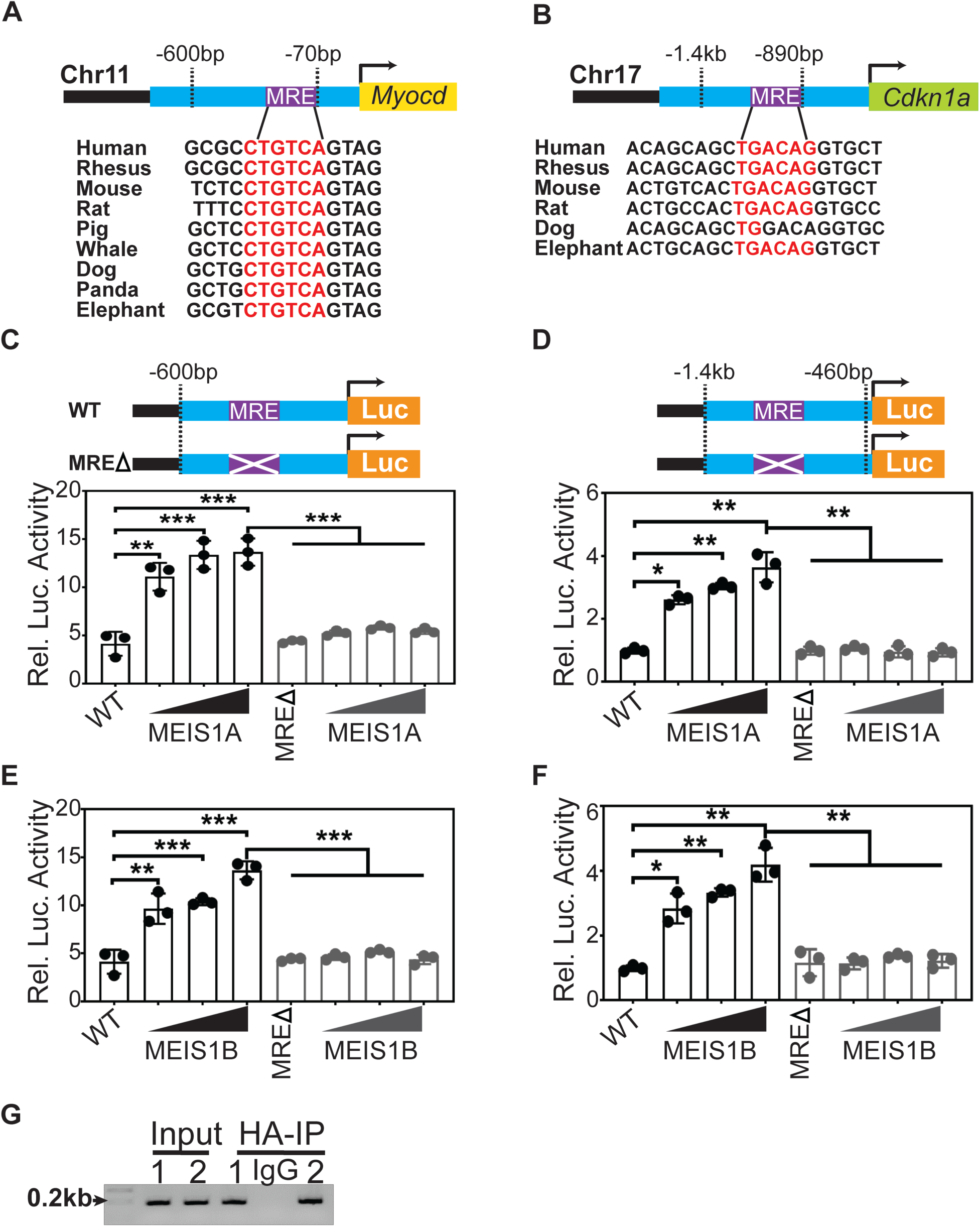
Regulation of *Myocd* by MEIS1 in VSMC. **A-B)** schematic showing the conservation of MRE in the upstream promotor of *Myocd* in (A) and *Cdkn1a* in (B). **C**-**D**) Luciferase assay shows the activation of *Myocd* and *Cdkn1a* prmotoers with increasing *MEIS1A*, *MEIS1B* isoforms with WT, but not MRE mutant. Purple X= MREΔ

## 4. Discussion

In this study, we identified MEIS1 as a novel, SRF/MYOCD-independent transcriptional regulator essential for maintaining VSMC differentiation. Although the MEIS1 promoter contains conserved CArG motifs, perturbation of SRF or MYOCD had no effect on MEIS1 mRNA levels. Instead, MEIS1 is robustly expressed in quiescent VSMCs and is rapidly downregulated during dedifferentiation; in human atherosclerotic plaques, passaged VSMCs in vitro, and multiple rodent injury models concomitant with the loss of canonical contractile markers (LMOD1, MYH11). Functionally, viral-mediated MEIS1 knockdown drives VSMC phenotypic switching, evidenced by decreased CDKN1A and contractile gene expression alongside increased KI67 and KLF4, whereas adenoviral MEIS1 overexpression elevates cell-cycle inhibitors and contractile proteins.

CRISPR-Cas9–mediated deletion of endogenous *Meis1* in MOVAS cells recapitulated the dedifferentiated, hyperproliferative phenotype and confirmed MEIS1’s necessity for contractile gene maintenance. Mechanistically, we discovered a conserved MEIS1 response element (MRE) on the antisense strand 75 bp upstream of the Myocd transcription start site, an unconventional arrangement in mammals, through which MEIS1 directly binds and activates Myocd and *Cdkn1a* promoter.

Our results extend prior observations of MEIS1 occupancy at the Cdkn1a locus and its role in pulmonary artery smooth muscle cell^51^ by providing comprehensive gain- and loss-of-function analyses and the first true *Meis1* null VSMC line to validate antibody specificity and enable future *in vivo* studies. Nonetheless, our work is limited by reliance on mRNA and RNA-scope in lieu of a specific MEIS1 antibody, potential off-target or supra-physiological effects of shRNA and viral overexpression, lack of conditional, VSMC-specific *Meis1* knockout mice for definitive *in vivo* validation, and absence of genome-wide ChIP-Seq to map MEIS1’s full regulatory network. Future efforts employing conditional knockout models, temporal single-cell analyses, and chromatin profiling will be critical to fully elucidate MEIS1’s role in vascular disease and explore its therapeutic potential.

Importantly, we also generated and validated a conditional *HAMeis1* allele that yields complete loss of MEIS1 protein following Cre-mediated recombination. This model provides a true conditional null allele, enables reliable detection of endogenous MEIS1 protein through the HA tag, and overcomes the incomplete knockout associated with the previously described exon 8-targeted floxed allele. Nonetheless, our work remains limited by potential supra-physiological effects of shRNA and viral overexpression, the absence of an *in vivo* VSMC-specific phenotypic analysis using the newly generated conditional allele, and the absence of genome-wide ChIP-seq to define MEIS1’s broader regulatory network. Future studies employing VSMC-specific deletion of the validated Meis1 allele, temporal single-cell analyses, and chromatin profiling will be important for defining the contribution of MEIS1 to VSMC phenotypic modulation and vascular disease.

## Acknowledgments/Funding

Research reported in this publication was supported by the National Institutes of Health under Award Number K99HL169827 to A.R.S. Additional support was provided by LSU Health Shreveport startup funds. We also acknowledge the LSU Health Shreveport Research Core Facility (RRID:SCR_024775) for research support and core services.

## Author Contributions

AW and ARS designed and performed the experiments. SJ contributed to experimental studies and data analysis. AK assisted with experimental design and troubleshooting. OJS performed histological sectioning. JD and SHG managed mouse breeding, genotyping, tissue isolation, and staining. KD performed the snATAC-seq and ChIP-seq analyses. LM and UH provided the BiKE and BiRCA datasets and contributed to their interpretation. GC provided human carotid artery samples. XL and JMM contributed to study design and supervision. ARS conceived and supervised the study, acquired funding, and drafted the manuscript. All authors reviewed and approved the final manuscript.

**Figure S1. *MEIS1* expression is MYOCD/SRF/CArG independent. A)** SRF-Chip-seq in Human Coronary Artery Smooth Muscle (HCASM) and endothelial cells (HUVEC) showed an SRF binding over a conserved CArG box in the up-stream promotor of *MEIS1*, and luciferase assay in A7r5 cells showed a mild induction of SRF to *Mesi1* promoter. **B-C**) qRT-PCR of *MEIS1*, *MEIS2*, *MEIS3* in HCASMC knockdown of MYOCD in (**C**) and over expression in (**D**). **D-E**) qRT-PCR of *MEIS1*, *MEIS2*, *MEIS2* in SRF-knockdown in two different HCASMC Lonza in (**D**), and Promo in (**E**). Each dot is represents the mean of three technical replicates.

**Figure S2. *MEIS1* expression is MYOCD/SRF/CArG independent. A)** qRT-PCR of *TAGLN*, *MEIS1*, *MEIS2*, *MEIS3* in SRF-knockdown in HCASMC Thermo. **B**) Western blot of SRF in HCASMC Thermo and PROMO. **C-G)** Reads counts of *SRF*, *TAGLN*, *CNN1*, *MYOCD*, and *MEIS1* from RNA-seq in shSRF in three different HCAMSC Lonza, Promo, and thermo. Each dot is a representative for a biological repeat

**Figure S3. Tissue expression profile of *MEIS1*, *MEIS2*, and *MEIS3*. A)** Screenshot of GTEx portal data (https://www.gtexportal.org/home/) for *MEIS1* in (**A**), *MEIS2* in (**B**), and *MEIS3* in (**C**). **D-F**) qRT-PCR expression of *Meis1* (**D**), *Meis2* (**E**), and *Meis3* (**F**) in different mouse tissues (Male and females). Red rectangle highlighted the expression in aortae. The experimental is a representative for at least three biological repeats.

**Figure S4. Ectopic *in vivo* delivery of adenovirus. A)** Experimental design [day 0-left carotid artery (LCA) wrapped with pluronic gel containing adenovirus (GFP as control or Ad-shMEIS1-GFP), Day 14, LCA was isolated for qRT-PCR, Western blot, and Immunofluorescence]. **B**) Immunostaining of GFP showed successful delivery of the adenovirus to the medial layer of the carotid artery. N.C is a negative control.

**Figure S5. Validation of commercial MEIS1 antibodies in null MEIS1 MOVAS cells. A)** Three commercial Anti-MEIS1 antibodies from abcam (cat# ab124686, cat# ab19867, cat# ab229962) and **B**) anti-MEIS1 antibody from Millipore (cat# ABE2864). WT= wild type, and KO= knockout clones,1,2,3 are individual MOVAS-single cell clones,

## Notes

### Competing Interest Statement

The authors have declared no competing interest.

## References

1. Mack CP. Signaling mechanisms that regulate smooth muscle cell differentiation. *Arteriosclerosis*, Thrombosis and Vascular Biology. 2011;31:1495–1505.

2. Owens GK, Kumar MS, Wamhoff BR. Molecular regulation of vascular smooth muscle cell differentiation in development and disease. Physiological Reviews. 2004;84:767–801.

3. Khachigian LM, Black BL, Ferdinandy P, De Caterina R, Madonna R, Geng YJ. Transcriptional regulation of vascular smooth muscle cell proliferation, differentiation and senescence: Novel targets for therapy. Vascul Pharmacol. 2022;146:107091. doi: 10.1016/j.vph.2022.107091

4. Norman C, Runswick M, Pollock R, Treisman R. Isolation and properties of cDNA clones encoding SRF, a transcription factor that binds to the c-fos serum response element. Cell. 1988;55:989–1003.

5. Minty A, Kedes L. Upstream regions of the human cardiac actin gene that modulate its transcription in muscle cells: presence of an evolutionarily conserved repeated motif. Molecular and Cellular Biology. 1986;6:2125–2136.

6. Miano JM. Serum response factor: toggling between disparate programs of gene expression. Journal of Molecular Cellular Cardiology. 2003;35:577–593.

7. Onuh JO, Qiu H. Serum response factor-cofactor interactions and their implications in disease. FEBS J. 2021;288:3120–3134. doi: 10.1111/febs.15544

8. Miano JM, Long X, Fujiwara K. Serum response factor: master regulator of the actin cytoskeleton and contractile apparatus. American Journal of Physiology: Cell Physiology. 2007;292:C70–C81.

9. Wang DZ, Chang PS, Wang Z, Sutherland L, Richardson JA, Small E, Krieg PA, Olson EN. Activation of cardiac gene expression by myocardin, a transcriptional cofactor for serum response factor. Cell. 2001;105:851–862.

10. Chen J, Kitchen CM, Streb JW, Miano JM. Myocardin: a component of a molecular switch for smooth muscle differentiation. J Mol Cell Cardiol. 2002;34:1345–1356. doi: 10.1006/jmcc.2002.2086

11. Du K, Ip HS, Li J, Chen M, Dandre F, Yu W, Lu MM, Owens GK, Parmacek MS. Myocardin is a critical serum response factor cofactor in the transcriptional program regulating smooth muscle cell differentiation. Molecular and Cellular Biology. 2003;23:2425–2437.

12. Yoshida T, Sinha S, Dandre F, Wamhoff BR, Hoofnagle MH, Kremer BE, Wang DZ, Olson EN, Owens GK. Myocardin is a key regulator of CArG-dependent transcription of multiple smooth muscle marker genes. Circulation Research. 2003;92:856–864.

13. Wang Z, Wang DZ, Pipes GCT, Olson EN. Myocardin is a master regulator of smooth muscle gene expression. Proceedings of the National Academy of Sciences of the United States of America. 2003;100:7129–7134.

14. Espinosa-Diez C, Mandi V, Du M, Liu M, Gomez D. Smooth muscle cells in atherosclerosis: clones but not carbon copies. JVS Vasc Sci. 2021;2:136–148. doi: 10.1016/j.jvssci.2021.02.002

15. Worssam MD, Jorgensen HF. Mechanisms of vascular smooth muscle cell investment and phenotypic diversification in vascular diseases. Biochem Soc Trans. 2021;49:2101–2111. doi: 10.1042/BST20210138

16. Wirka RC, Wagh D, Paik DT, Pjanic M, Nguyen T, Miller CL, Kundu R, Nagao M, Coller J, Koyano TK, et al. Atheroprotective roles of smooth muscle cell phenotypic modulation and the TCF21 disease gene as revealed by single-cell analysis. Nat Med. 2019;25:1280–1289. doi: 10.1038/s41591-019-0512-5

17. Alencar GF, Owsiany KM, K S, Sukhavasi K, Mocci G, Nguyen A, Williams CM, Shamsuzzaman S, Mokry M, Henderson CA, et al. The stem cell pluripotency genes Klf4 and Oct4 regulate complex SMC phenotypic changes critical in late-stage atherosclerotic lesion pathogenesis. Circulation. 2020;142:2045–2059. doi: 10.1161/CIRCULATIONAHA.120.046672

18. Pan H, Xue C, Auerbach BJ, Fan J, Bashore AC, Cui J, Yang DY, Trignano SB, Liu W, Shi J, et al. Single-Cell Genomics Reveals a Novel Cell State During Smooth Muscle Cell Phenotypic Switching and Potential Therapeutic Targets for Atherosclerosis in Mouse and Human. Circulation. 2020;142:2060–2075. doi: 10.1161/CIRCULATIONAHA.120.048378

19. Tharp DL, Wamhoff BR, Turk JR, Bowles DK. Upregulation of intermediate-conductance Ca2+-activated K+ channel (IKCa1) mediates phenotypic modulation of coronary smooth muscle. American Journal of Physiology: Heart and Circulatory Physiology. 2006;291:H2493–H2503.

20. Ackers-Johnson M, Talasila A, Sage AP, Long X, Bot I, Morrell NW, Bennett MR, Miano JM, Sinha S. Myocardin regulates vascular smooth muscle cell inflammatory activation and disease. Arterioscler Thromb Vasc Biol. 2015;35:817–828. doi: 10.1161/ATVBAHA.114.305218

21. Long X, Tharp DL, Georger MA, Slivano OJ, Lee MY, Wamhoff BR, Bowles DK, Miano JM. The smooth muscle cell-restricted KCNMB1 ion channel subunit is a direct transcriptional target of serum response factor and myocardin. Journal of Biological Chemistry. 2009;284:33671–33682.

22. Minami T, Kuwahara K, Nakagawa Y, Takaoka M, Kinoshita H, Nakao K, Kuwabara Y, Yamada Y, Yamada C, Shibata J, et al. Reciprocal expression of MRTF-A and myocardin is crucial for pathological vascular remodelling in mice. EMBO Journal. 2012;31:4428–4440.

23. Wang Z, Wang DZ, Hockemeyer D, McAnally J, Nordheim A, Olson EN. Myocardin and ternary complex factors compete for SRF to control smooth muscle gene expression. Nature. 2004;428:185–189. doi: 10.1038/nature02382

24. Tang Rh, Zheng XL, Callis TE, Stansfield WE, He J, Baldwin AS, Wang DZ, Selzman CH. Myocardin inhibits cellular proliferation by inhibiting NF-k B(p65)-dependent cell cycle progression. Proceedings of the National Academy of Sciences of the United States of America. 2008;105:3362–3367.

25. Xu Z, Ji G, Shen J, Wang X, Zhou J, Li L. SOX9 and myocardin counteract each other in regulating vascular smooth muscle cell differentiation. Biochemical and biophysical research communications. 2012;422:285–290. doi: 10.1016/j.bbrc.2012.04.149

26. Liu F, Wang X, Hu G, Wang Y, Zhou J. The transcription factor TEAD1 represses smooth muscle-specific gene expression by abolishing myocardin function. Journal of Biological Chemistry. 2014;289:3308–3316. doi: 10.1074/jbc.M113.515817

27. Zheng JP, He X, Liu F, Yin S, Wu S, Yang M, Zhao J, Dai X, Jiang H, Yu L, et al. YY1 directly interacts with myocardin to repress the triad myocardin/SRF/CArG box-mediated smooth muscle gene transcription during smooth muscle phenotypic modulation. Sci Rep. 2020;10:21781. doi: 10.1038/s41598-020-78544-3

28. Miano JM, Fisher EA, Majesky MW. Fate and State of Vascular Smooth Muscle Cells in Atherosclerosis. Circulation. 2021;143:2110–2116. doi: 10.1161/CIRCULATIONAHA.120.049922

29. Ueyama T, Kasahara H, Ishiwata T, Nie Q, Izumo S. Myocardin expression is regulated by Nkx2.5, and its function is required for cardiomyogenesis. Molecular and Cellular Biology. 2003;23:9222–9232.

30. Xie WB, Li Z, Miano JM, Long X, Chen SY. Smad3-mediated myocardin silencing: a novel mechanism governing the initiation of smooth muscle differentiation. Journal of Biological Chemistry. 2011;286:15050–15057.

31. Shi N, Chen SY. Cell division cycle 7 mediates transforming growth factor-beta-induced smooth muscle maturation through activation of myocardin gene transcription. Journal of biological chemistry. 2013;288:34336–34342. doi: 10.1074/jbc.M113.498238

32. Wang L, Qiu P, Jiao J, Hirai H, Xiong W, Zhang J, Zhu T, Ma PX, Chen YE, Yang B. Yes-associated protein inhibits transcription of myocardin and attenuates differentiation of vascular smooth muscle cell from cardiovascular progenitor cell lineage. Stem Cells. 2016. doi: 10.1002/stem.2484

33. Lei X, Zhao J, Sagendorf JM, Rajashekar N, Xu J, Dantas Machado AC, Sen C, Rohs R, Feng P, Chen L. Crystal structures of ternary complexes of MEF2 and NKX2-5 bound to DNA reveal a disease related protein-protein interaction interface. J Mol Biol. 2020;432:5499–5508. doi: 10.1016/j.jmb.2020.07.004

34. Tan Z, Li J, Zhang X, Yang X, Zhang Z, Yin KJ, Huang H. P53 promotes retinoid acid-induced smooth muscle cell differentiation by targeting myocardin. Stem Cells Dev. 2018;27:534–544. doi: 10.1089/scd.2017.0244

35. Yang X, Gong Y, Tang Y, Li H, He Q, Gower L, Liaw L, Friesel RE. Spry1 and Spry4 differentially regulate human aortic smooth muscle cell phenotype via Akt/FoxO/myocardin signaling. PLoS One. 2013;8:e58746. doi: 10.1371/journal.pone.0058746

36. Jin Y, Xie Y, Ostriker AC, Zhang X, Liu R, Lee MY, Leslie KL, Tang W, Du J, Lee SH, et al. Opposing actions of AKT (Protein Kinase B) isoforms in vascular smooth muscle injury and therapeutic response. Arterioscler Thromb Vasc Biol. 2017;37:2311–2321. doi: 10.1161/ATVBAHA.117.310053

37. Shikatani EA, Chandy M, Besla R, Li CC, Momen A, El-Mounayri O, Robbins CS, Husain M. c-Myb regulates proliferation and differentiation of adventitial Sca1+ vascular smooth muscle cell progenitors by transactivation of myocardin. Arterioscler Thromb Vasc Biol. 2016;36:1367–1376. doi: 10.1161/ATVBAHA.115.307116

38. Liu R, Jin Y, Tang WH, Qin L, Zhang X, Tellides G, Hwa J, Yu J, Martin KA. Ten-eleven translocation-2 (TET2) is a master regulator of smooth muscle cell plasticity. Circulation. 2013;128:2047–2057. doi: 10.1161/CIRCULATIONAHA.113.002887

39. Turner EC, Huang CL, Govindarajan K, Caplice NM. Identification of a Klf4-dependent upstream repressor region mediating transcriptional regulation of the myocardin gene in human smooth muscle cells. Biochimica et biophysica acta. 2013;1829:1191–1201. doi: 10.1016/j.bbagrm.2013.09.002

40. Huang Y, Lin L, Yu X, Wen G, Pu X, Zhao H, Fang C, Zhu J, Ye S, Zhang L, et al. Functional involvement of heterogeneous nuclear ribonucleoprotein A1 in smooth muscle differentiation from stem cells in vitro and in vivo. Stem Cells. 2013;31:906–917. doi: 10.1002/stem.1324

41. Wang K, Long B, Zhou J, Li PF. miR-9 and NFATc3 regulate myocardin in cardiac hypertrophy. Journal of Biological Chemistry. 2010;285:11903–11912.

42. Creemers EE, Sutherland LB, McNally J, Richardson JA, Olson EN. Myocardin is a direct transcriptional target of Mef2, Tead and Foxo proteins during cardiovascular development. Development. 2006;133:4245–4256.

43. Sun Q, Chen G, Streb JW, Long X, Yang Y, Stoeckert CJ, Jr., Miano JM. Defining the mammalian CArGome. Genome Research. 2006;16:197–207.

44. Caronia JM, Sorensen DW, Leslie HM, van Berlo JH, Azarin SM. Adhesive thermosensitive gels for local delivery of viral vectors. Biotechnol Bioeng. 2019;116:2353–2363. doi: 10.1002/bit.27007

45. Lyu Q, Dhagia V, Han Y, Guo B, Wines-Samuelson ME, Christie CK, Yin Q, Slivano OJ, Herring P, Long X, et al. CRISPR-Cas9 mediated epitope tagging provides accurate and versatile assessment of myocardin. Arterioscler Thromb Vasc Biol. 2018;38:2184–2190. doi: 10.1161/ATVBAHA.118.311171

46. Curaj A, Zhoujun W, Staudt M, Liehn EA. Induction of Accelerated Atherosclerosis in Mice: The “Wire-Injury” Model. J Vis Exp. 2020. doi: 10.3791/54571

47. Takayama T, Shi X, Wang B, Franco S, Zhou Y, DiRenzo D, Kent A, Hartig P, Zent J, Guo LW. A murine model of arterial restenosis: technical aspects of femoral wire injury. J Vis Exp. 2015. doi: 10.3791/52561

48. Perisic L, Aldi S, Sun Y, Folkersen L, Razuvaev A, Roy J, Lengquist M, Åkesson S, Wheelock CE, Maegdefessel L, et al. Gene expression signatures, pathways and networks in carotid atherosclerosis. Journal of Internal Medicine. 2016;279:293–308. doi: 10.1111/joim.12448

49. Wadén K, Hultgren R, Kotopouli MI, Gillgren P, Roy J, Hedin U, Matic L. Long Term Mortality Rate in Patients Treated with Carotid Endarterectomy. European Journal of Vascular and Endovascular Surgery. 2023;65:778–786. doi: 10.1016/j.ejvs.2023.02.079

50. Röhl S, Rykaczewska U, Seime T, Suur BE, Diez MG, Gådin JR, Gainullina A, Sergushichev AA, Wirka R, Lengquist M, et al. Transcriptomic profiling of experimental arterial injury reveals new mechanisms and temporal dynamics in vascular healing response. JVS-Vascular Science. 2020;1:13–27. doi: 10.1016/j.jvssci.2020.01.001

51. Yao MZ, Ge XY, Liu T, Huang N, Liu H, Chen Y, Zhang Z, Hu CP. MEIS1 regulated proliferation and migration of pulmonary artery smooth muscle cells in hypoxia-induced pulmonary hypertension. Life Sci. 2020;255:117822. doi: 10.1016/j.lfs.2020.117822

52. Turner AW, Hu SS, Mosquera JV, Ma WF, Hodonsky CJ, Wong D, Auguste G, Song Y, Sol-Church K, Farber E, et al. Single-nucleus chromatin accessibility profiling highlights regulatory mechanisms of coronary artery disease risk. Nature Genetics. 2022;54:804–816. doi: 10.1038/s41588-022-01069-0

53. Doudna JA, Charpentier E. Genome editing. The new frontier of genome engineering with CRISPR-Cas9. Science. 2014;346:1258096. doi: 10.1126/science.1258096

54. Miano JM, Zhu QM, Lowenstein CJ. A CRISPR path to engineering new genetic mouse models for cardiovascular research. *Arteriosclerosis*, Thrombosis and Vascular Biology. 2016;36:1058–1075.

55. Afroze T, Yang LL, Wang C, Gros R, Kalair W, Hoque AN, Mungrue IN, Zhu Z, Husain M. Calcineurin-independent regulation of plasma membrane Ca2+ ATPase-4 in the vascular smooth muscle cell cycle. American Journal of Physiology Cell Physiology. 2003;285:C88–C95.

56. Castro-Mondragon JA, Riudavets-Puig R, Rauluseviciute I, Lemma RB, Turchi L, Blanc-Mathieu R, Lucas J, Boddie P, Khan A, Manosalva Perez N, et al. JASPAR 2022: the 9th release of the open-access database of transcription factor binding profiles. Nucleic Acids Res. 2022;50:D165–D173. doi: 10.1093/nar/gkab1113

