## Supplementary Figures for "Myeloid Ecotropic Viral Integration Site 1 Promotes Vascular Smooth Muscle Differentiation through Activation of Myocardin"

**A**

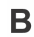

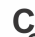

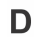

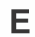

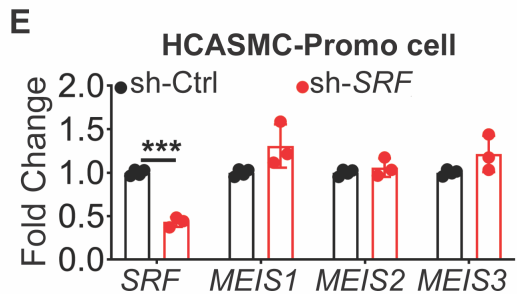

(Wally et al, supplementary Fig2)

**A**

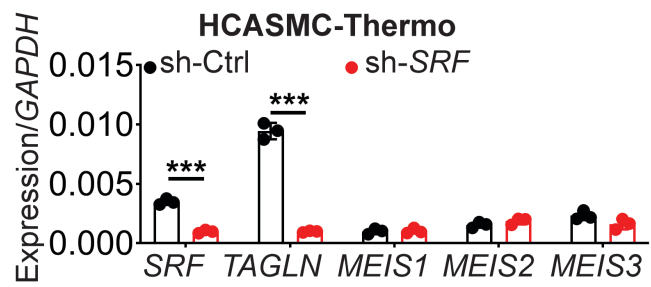

**B**

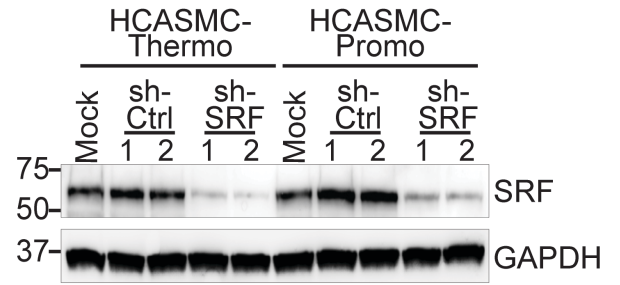

**C**

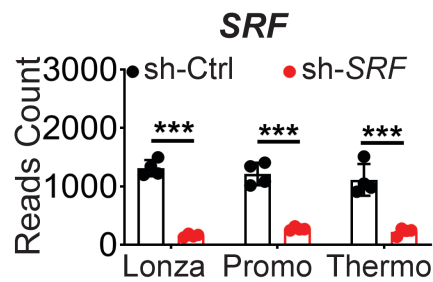

**D**

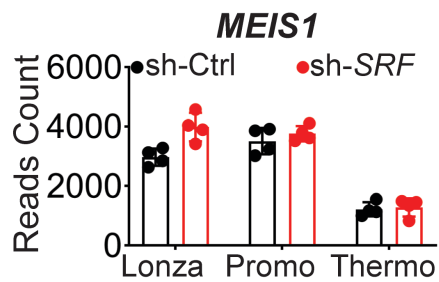

**E**

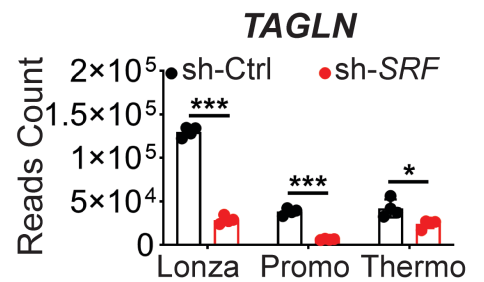

**F**

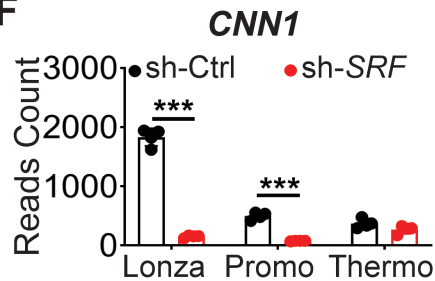

**G**

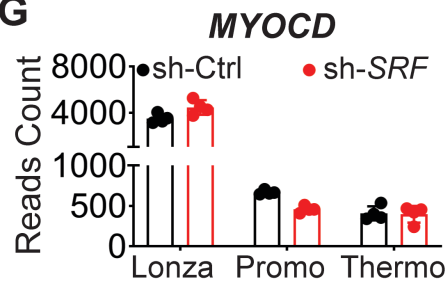

(Wally et al, supplementary Fig3)

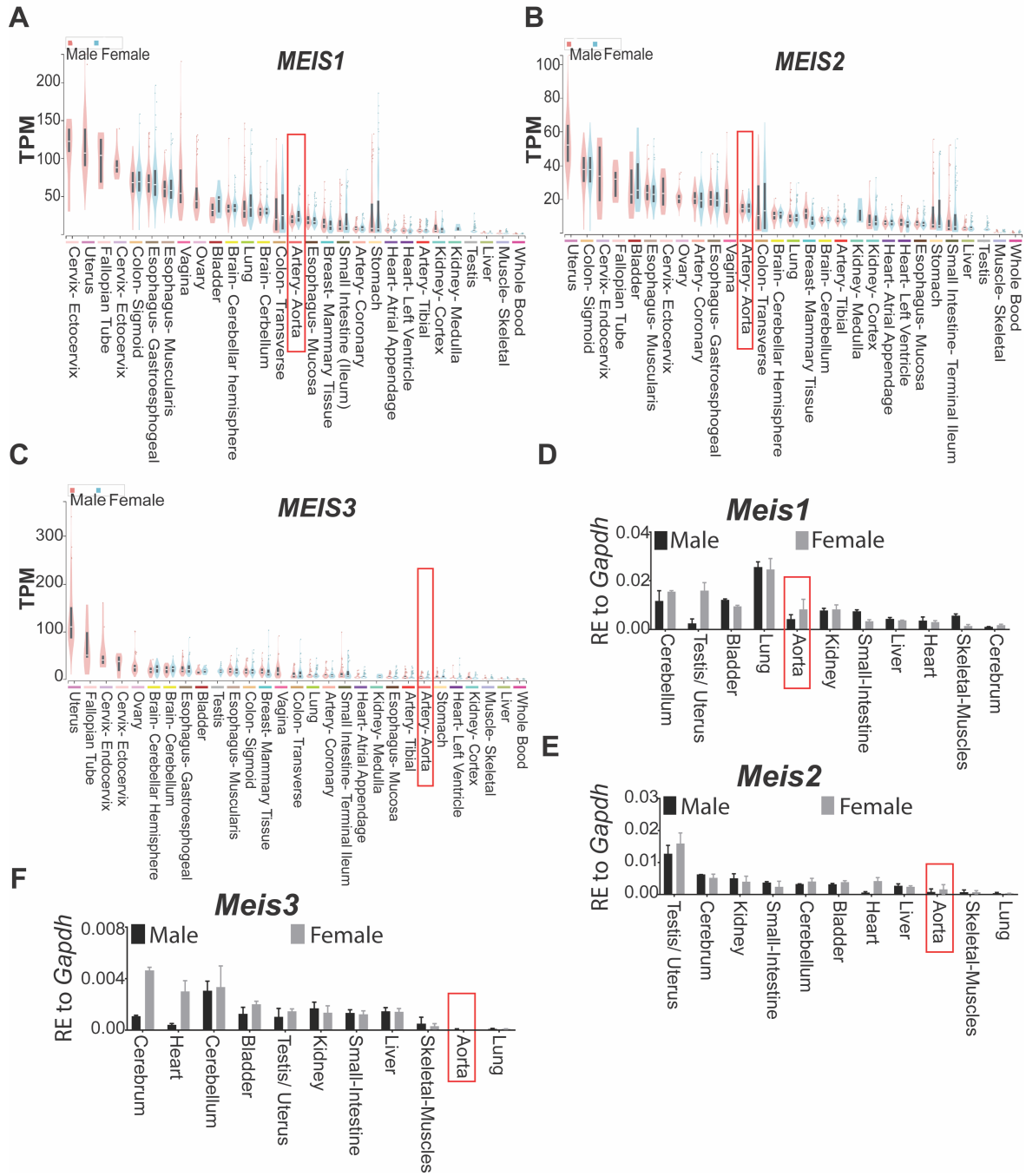

**A**

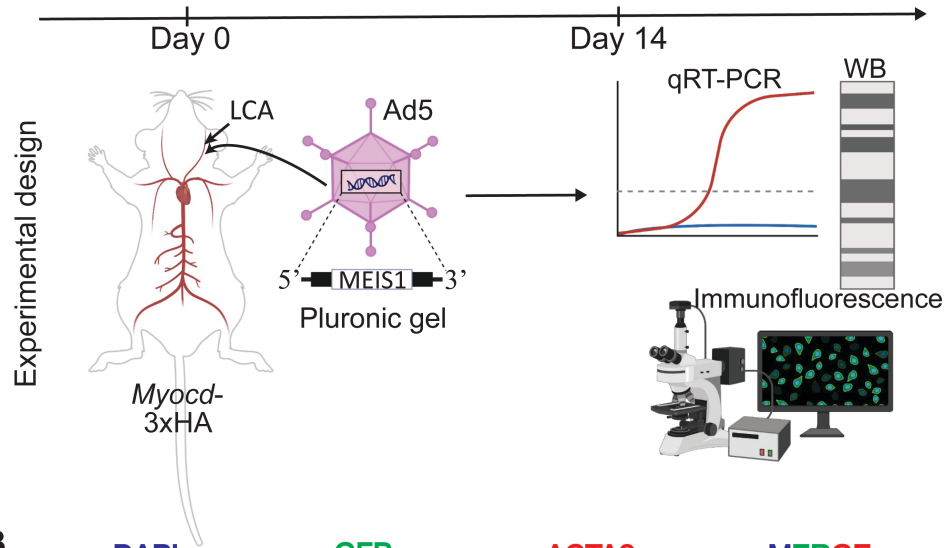

**B**

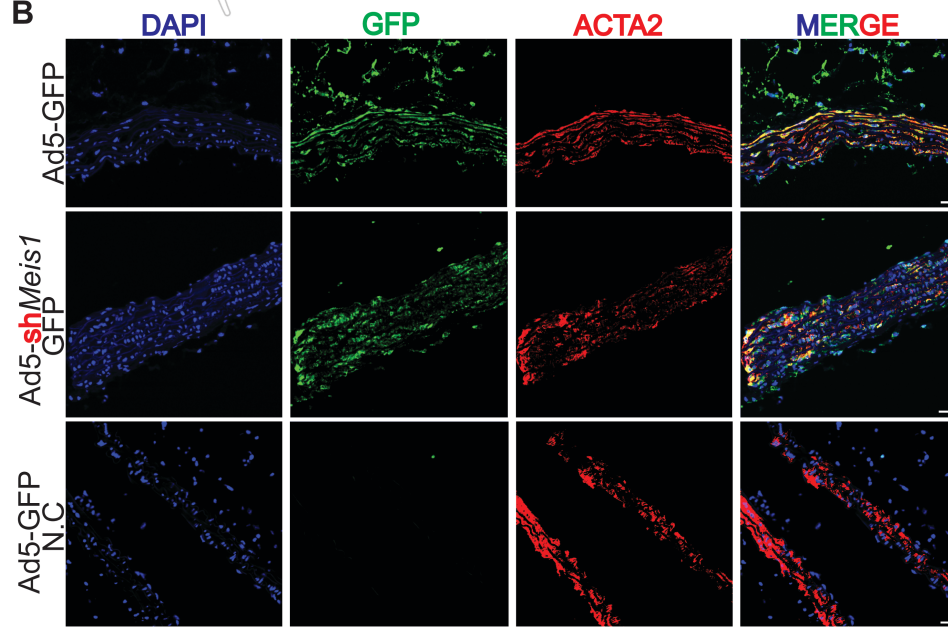

(Wally et al, supplementary Fig5)

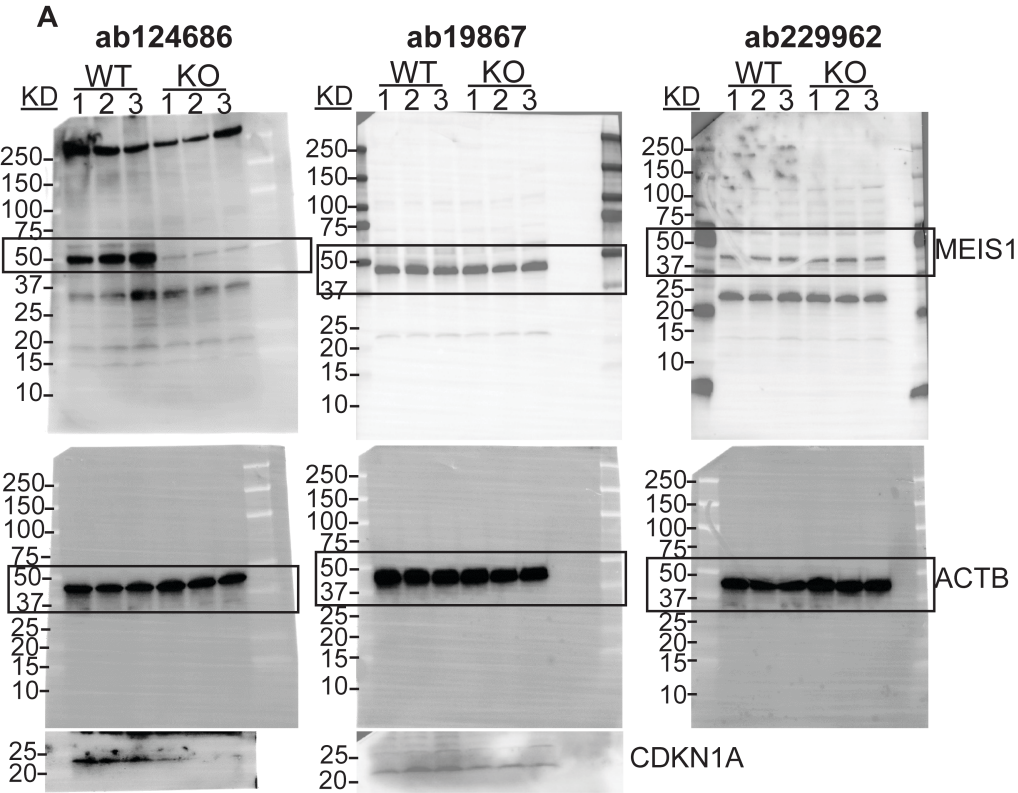

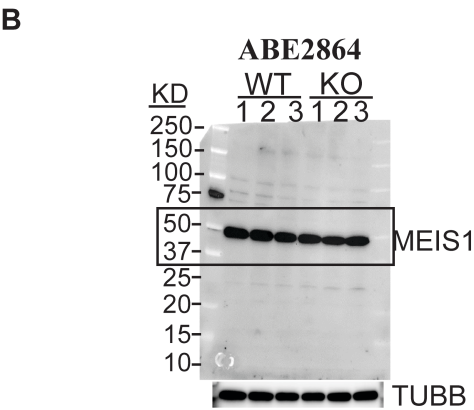
